# ARGeement: sequence-based, versioned mapping of antimicrobial resistance gene databases

**DOI:** 10.64898/2026.11.17.767227

**Authors:** Vesna Vujić, Georgina Helyes, Adrienn Gréta Tóth, István Csabai, Norbert Solymosi

## Abstract

Genomic studies of antimicrobial resistance (AMR) rely on several independently curated reference databases, which describe the same genes under different names, in different structures and with different classifications. This makes results obtained with different databases difficult to compare. ARGeement maps the entries of AMRFinderPlus, the Comprehensive Antibiotic Resistance Database (CARD) and ResFinder/DisinFinder to each other by protein sequence, independently of their names. It characterises every link by its relation type, difference count and coverage, tracks the database versions, and produces a documented, immutable mapping for every release. The 17,920 entries of the release described here encode 17,959 protein sequences, corresponding to 9,337 unique proteins. For 64.4% of the cross-database entry pairs that encode the same protein, the names differ even when letter case is ignored, and 447 pairs show a genuine naming conflict. Conversely, 2.0% of the pairs with the same allele name do not encode the same protein. For synthetic variants created with known modifications, the original protein was among the best hits in 99.6% of the cases. In 100 complete RefSeq genomes annotated with AMRFinderPlus, the Resistance Gene Identifier (RGI) and ResFinder, the gene names agreed at 55.3%–75.6% of the loci called by both tools of a pair, whereas after mapping the two reference proteins were identical, near-identical or length variants at 89.6%–94.2%. These results show that name-based comparison is unreliable even for identical sequences, and that a sequence-based, versioned mapping is needed.

## 1 Introduction

Whole-genome and metagenome sequencing have become essential tools for studying antimicrobial resistance (AMR). The identification of antimicrobial resistance genes (ARGs) relies on reference databases, of which the most widely used are the NCBI AMRFinderPlus database [1], the Comprehensive Antibiotic Resistance Database (CARD) [2] and ResFinder [3]. They are curated by different institutions, with different aims and structures. AMRFinderPlus is built on a curated gene hierarchy and allele-level nomenclature, CARD on an ontology (the Antibiotic Resistance Ontology, ARO) and a system of detection models, and ResFinder on nucleotide references. DisinFinder, a companion database of ResFinder, contains disinfectant and biocide resistance genes [4].

The results of sequencing-based AMR studies depend strongly on the chosen reference database and annotation tool [5]. In an international inter-laboratory study, the participating laboratories obtained different resistance predictions from the whole-genome data of the same isolates, and a substantial part of the discrepancies was attributable to differences between the databases and the tools [6]. A systematic review likewise found considerable differences in the content of resistance marker databases and in the accuracy of phenotype prediction [7]. The EUCAST subcommittee therefore highlights the quality and maintenance of reference databases among the conditions for genome-based susceptibility prediction [8]. Genomic AMR surveillance yields comparable results only if the gene calls obtained with different databases can be mapped onto each other.

In an earlier review [9], we compared the ARG databases available at the time and found, among other things, that their nomenclature and classification differ substantially, which hampers the comparison of results from different sources. We proposed the development of a standardisation procedure. Since then, the field has changed: several databases are no longer updated, or only rarely, and the secondary databases that are still maintained to some extent (e.g. MEGARes [10]) are built largely from the primary sources above. Several approaches exist for harmonising results and databases. One is to unify the output format of the annotation tools. hAMRonization, based on the PHA4GE specification, converts the output of 18 tools to a common structure, but leaves the gene names as given by the tools [11]. Another is to map the annotations onto an ontology: argNorm normalises the output of annotation tools to the CARD ontology (ARO) using mapping tables prepared in advance for fixed database versions [12]. A third is to build secondary databases that merge the sources under their own nomenclature and hierarchy, for example ARG-ANNOT [13], MEGARes [10], SARG [14], DeepARG-DB [15] and PanRes [16], which also offers structure-based homology search. To our knowledge, none of the published approaches addresses how the entries of two databases relate to each other *by sequence*, and how this relationship changes as the databases are updated.

Our aim was to develop a tool (ARGeement) that (i) maps ARG entries to each other by protein sequence, independently of their names, (ii) characterises the links quantitatively (identity, difference count, coverage), (iii) tracks the database versions automatically while retaining the mappings built for earlier versions, and (iv) produces a documented, publishable result for every database release.

## 2 Methods

### 2.1 Overview of the tool

Throughout, the term antimicrobial resistance gene (ARG) also covers disinfectant and biocide resistance genes. Three primary, continuously maintained databases were included in the comparison (Table 1). The entries of DisinFinder, the companion database of ResFinder, were used in the mapping, whereas the PointFinder [17] point mutation database was used only to describe the content of the databases (Section 2.7). Each reference sequence in a database that describes an ARG or one of its alleles is called an *ARG entry*. A gene may have several ARG entries if the database lists its alleles separately. Because of ORF splitting (Section 2.2), one ARG entry may give rise to several proteins. We therefore distinguish the entries of the source database from *protein-level entries*. The latter are the entries translated as a whole and the retained ORF sub-entries obtained by splitting. AMRFinderPlus and CARD store protein sequences, so every included entry of these is also a protein-level entry.

**Table 1.** The included databases, the versions used and their numbers of entries. The entries of DisinFinder, the companion database of ResFinder, were used in the mapping, the PointFinder data only in the description of the mutation content of the databases. The Entries column gives the included source entries, the Protein-level column the protein-level entries (the entries translated as a whole and the retained ORF sub-entries), and the Unique proteins column the number of distinct protein sequences.

| Database | Version | Included content | Entries | Protein-level | Unique proteins |
|---|---|---|---|---|---|
| AMRFinderPlus | 2026-10-29.1 | catalogue type AMR (without point mutations, gene inactivations and susceptibility markers) and subtype STRESS/BIOCIDE | 8,633 | 8,633 | 8,633 |
| CARD | 4.0.2 | protein homolog models | 6,059 | 6,059 | 6,059 |
| ResFinder | 2.7.0 (git eb6c21a) | acquired resistance genes (nucleotide) | 3,212 | 3,249 | 3,162 |
| DisinFinder | 2.0.1 (git 09fc831) | disinfectant and biocide resistance genes (nucleotide) | 16 | 18 | 17 |
| PointFinder | 4.2.0 (git 3bd7d5e) | point mutations | – | – | – |

The processing workflow of ARGeement is shown in Fig. 1. The tool reads the database entries, translates the nucleotide sequences to protein, and compares every protein with every other. Based on the comparison it classifies the relations between entries, determines the families, harmonises the drug classes, and compares names with sequences in both directions. Mutation-based determinants are inventoried separately, by genus. The result is one release per profile. The workflow is run by Snakemake [18]; the individual steps are detailed in the following subsections.

**Figure 1.**
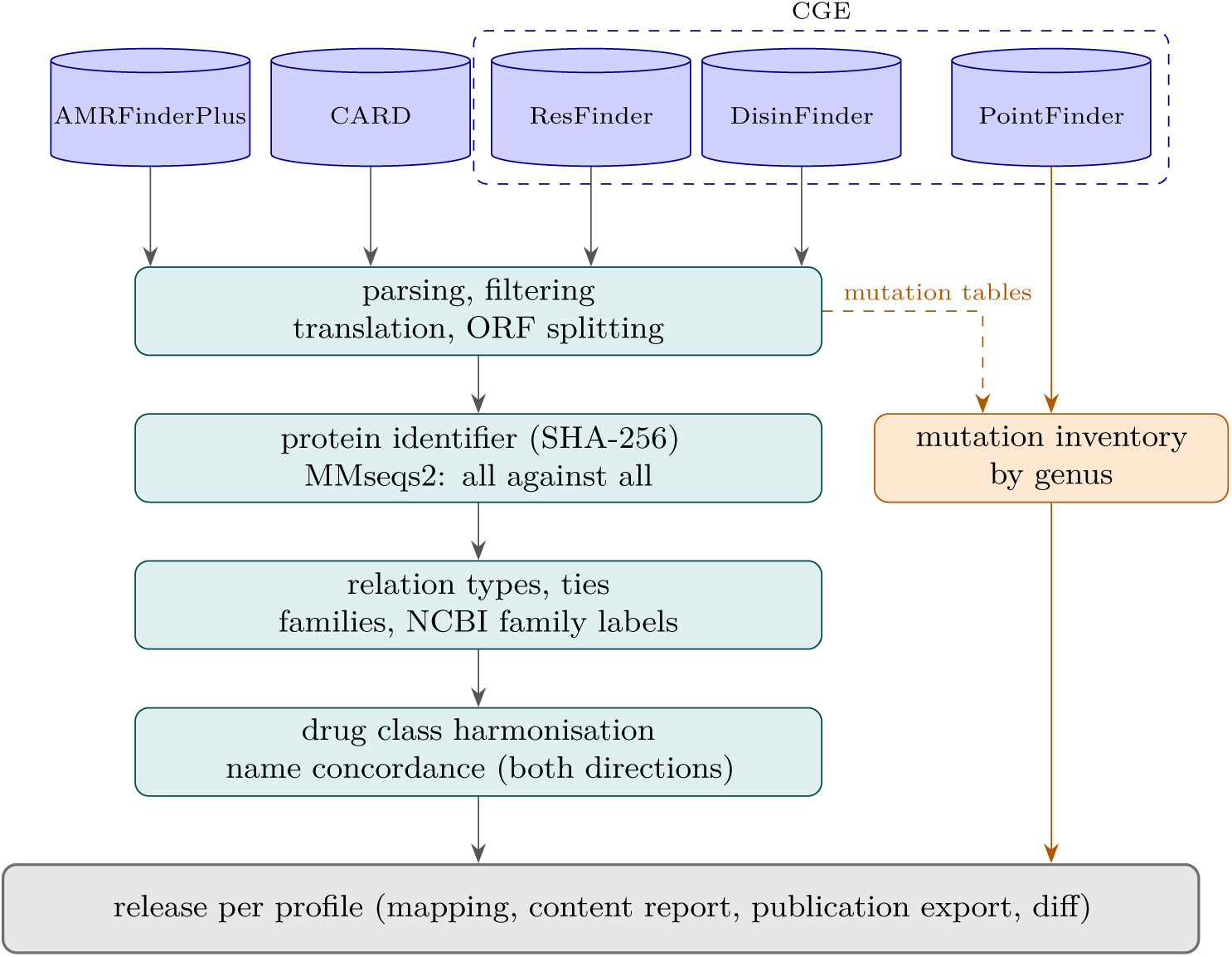
The processing workflow of ARGeement. ResFinder, DisinFinder and PointFinder are databases of the Center for Genomic Epidemiology (CGE). PointFinder and the mutation part of the databases enter only the mutation inventory and take no part in the sequence-based mapping.

### Profiles and releases

The tool is organised around *profiles*. A profile is a named set of settings that specifies which version of each database is used. The default profile always downloads the latest versions, whereas a pinned profile works from specified local database copies or specified git commits. If the profile also specifies an expected version, the tool checks that the version read matches it and stops if it does not. In this way a mapping can be built that matches exactly the results of an earlier ARG analysis performed with given database versions, without repeating the analysis. The result of a run of the tool is called a *release*. A release is a self-contained result package for a given profile and point in time. It contains the mapping of the entries, the content report and the publication export, as well as everything needed to reproduce the result, that is, the database versions, the settings and the tool versions used. Every profile has its own series of releases, and completed releases are never modified. A new release is built only if a new version of at least one database in the profile has appeared, so the series of releases follows the changes of the databases.

### Change tracking and publication

Comparing two releases (diff) reveals new, removed, renamed and sequence-changed entries, as well as changes in the relations. Manual review can thus be restricted to the items that actually changed. Like a relational database dump, the publication export consists of normalised, non-redundant tables (databases, relation types, families, proteins, entries and the relations of protein pairs), which reference each other through integer keys and from which the complete mapping can be restored. The tool is checked by automated tests on synthetic data, which run before every release is built; if a test fails, no release is built. The tests cover the inclusion filters, non-ATG start codons, genes stored on the reverse strand and in multigene operons, start codon ties, the exclusion of ORFs unrelated to resistance genes, a gene displaced by a longer overlapping ORF, the handling of ties and family labels, the exclusion of cluster chaining, the classification of drug classes and names, the restorability of the publication export, and the execution of the full workflow with two profiles and the comparison of the two releases.

### 2.2 Inclusion, translation and ORF splitting

The names and types of the AMRFinderPlus entries were taken from the Reference Gene Catalog. Rows of type AMR were included, except for the mutation- or context-dependent subtypes, that is, point mutations (POINT), gene inactivations (POINT_DISRUPT) and susceptibility markers (AMR-SUSCEPTIBLE). In addition, for comparison with DisinFinder, the BIOCIDE subtype of type STRESS was also included. The other stress response genes (metal, acid and heat stress) and the virulence genes were left out. Entries of both the core and the plus set were included, but the 3 entries flagged non-reported in the catalogue were not (Table 2). Several catalogue rows may belong to one protein accession (for example, a mutation row next to the gene’s own row for an acquired gene), so an entry was included if any of its rows met the criteria. Its name was the allele name of the first such row or, if missing, its gene family. Protein sequences were linked to the catalogue rows by the protein identifier in the FASTA header, accepting the GenBank identifier as well as the RefSeq identifier, so that entries without a RefSeq identifier (e.g. *norM-Vib*) were also included. Mutation data and proteins not listed in the catalogue were not used. The included and excluded categories of the source databases are summarised in Table 2.

**Table 2.** Entry categories of the source databases and their inclusion in the mapping. For AMRFinderPlus, the rows of the Reference Gene Catalog by type and subtype; for CARD, the model types. Of the included AMRFinderPlus categories, the core and plus rows were used; rows not reported by AMRFinderPlus (non-reported) are listed separately for each category. All ResFinder and DisinFinder entries were included.

| Database | Category | Count | Included |
|---|---|---|---|
| AMRFinderPlus | AMR/AMR | 8,569 | yes |
|  | AMR/POINT | 1,637 | no |
|  | VIRULENCE/VIRULENCE | 1,034 | no |
|  | STRESS/METAL | 182 | no |
|  | STRESS/BIOCIDE | 64 | yes |
|  | AMR/POINT_DISRUPT | 26 | no |
|  | AMR/AMR-SUSCEPTIBLE | 26 | no |
|  | STRESS/HEAT | 10 | no |
|  | STRESS/ACID | 3 | no |
|  | AMR/AMR (non-reported) | 3 | no |
| CARD | protein homolog model | 6,059 | yes |
|  | protein variant model | 239 | no |
|  | rRNA gene variant model | 86 | no |
|  | efflux pump system meta-model | 22 | no |
|  | protein knockout model | 16 | no |
|  | protein overexpression model | 15 | no |
|  | gene cluster meta-model | 14 | no |

From CARD, the protein sequences of the protein homolog models were used. The entry identifier is the ARO accession, and its name is the model name in the FASTA header (without the species name). If several sequences belonged to one ARO accession, they were distinguished by a serial number. The *AMR Gene Family*, drug classes and resistance mechanisms of the models were taken from the ontology categories in the database’s card.json file. Variant, overexpression, knockout and rRNA models describe resistance that depends on mutations or gene expression, and were therefore left out of the mapping. Their numbers by model type are given in Table 2, and the variant models were summarised by genus in the mutation comparison (Section 2.7).

The nucleotide sequences of ResFinder and DisinFinder were translated with the bacterial genetic code (NCBI translation table 11), with alternative start codons normalised to methionine. An entry was translated as a whole, as a single coding sequence, if its length is divisible by three, it contains no internal stop codon, and it starts with a valid start codon and ends with a stop codon. If the length is divisible by three and there is no internal stop codon but the start or stop codon is missing (truncated reference), the entry was also translated as a whole, unless an ORF on the reverse strand covers at least 80% of the sequence, in which case the gene was taken to be on the reverse strand. All other entries, such as operons containing several genes (e.g. *vanHAX*) and genes stored in a shifted frame, were split into open reading frames (ORFs) on both strands. All ORF candidates were identified in the six reading frames (ATG, GTG or TTG start codon, at least 60 amino acids). As these often overlap (the same gene may have several possible start codons), the non-overlapping combination covering the largest part of the sequence was selected. This is a weighted interval scheduling problem, which was solved exactly by dynamic programming [19]. To prevent a complete gene from being split into two shorter pieces merely because together they cover a few more nucleotides, 150 nucleotides were subtracted from the coverage of every selected ORF. ORFs starting with the rarer GTG and TTG start codons received a further deduction of 9.5 nucleotides. This rules out exact ties, and if a non-ATG start codon lies at most three codons upstream of an ATG, the shorter variant starting with ATG is selected. To allow for the short gene overlaps common in bacterial operons, adjacent ORFs were allowed to overlap by up to 8 nucleotides. A split entry does not count as a protein itself; only its ORFs take part in the mapping. This splitting may select ORFs other than the resistance gene, for example fragments of neighbouring genes included in the reference sequence or long ORFs on the opposite strand. Of the ORFs obtained by splitting, we therefore retained those that are identical to a curated (AMRFinderPlus or CARD) protein or are linked to one above the search lower bound. If none of the ORFs of an entry was linked to a curated protein, the splitting pattern decided. A single ORF covering at least 80% of the sequence was taken to be the gene itself, and several ORFs on the same strand that together covered at least 90% of the sequence were taken to be an operon absent from the curated databases (e.g. *sitABCD*); in both cases the ORFs were retained. If, however, at least one ORF was linked to a curated protein, the other ORFs were dropped. Excluded ORFs are flagged in the release but take no part in the mapping.

As the selection considers only the covered length, a longer, overlapping unrelated ORF could in principle displace the true gene. All ORF candidates of the split entries were therefore grouped by stop codon, the longest member of each group was searched against the curated proteins, and the selection was repeated so that candidates linked to a curated protein took precedence. The other candidates could only fill the remaining stretches.

### 2.3 Sequence comparison and relation types

Every protein received a unique identifier (the SHA-256 digest of its sequence), so entries encoding the same protein are linked without any threshold. Every unique protein was compared with every other unique protein using MMseqs2 [20] (minimum 50% sequence identity and 50% coverage). The search was run with the settings --min-seq-id 0.5 -c 0.5 --cov-mode 0 --seq-id-mode 0 -s 7.5 --max-seqs 20000 (MMseqs2 18.8cc5c). Sequence identity is relative to the alignment length, and coverage was computed separately for each protein. The maximum number of hits retained per query (--max-seqs) was raised from the default 300 to 20,000, because in large allele families (e.g. *blaPDC*, *blaOXA*) a protein has far more hits than this, and with a lower limit some of the close hits would also be lost. Every protein pair A–B was aligned in both directions (A to B and B to A), and the best alignment was kept for each direction. For every alignment we computed the *difference count* (*n*_diff_), that is, the number of mismatched and gap positions in the alignment plus the unaligned residues at the ends, as well as the coverage of both proteins (the length of the aligned stretch relative to the full length of the protein). At each end, only the longer of the two overhangs counts, so a substitution at the last position, which leaves one unaligned residue on each protein, counts as one difference, not two. The relation types and their classification thresholds are summarised in Table 3. The classification takes the coverage of both proteins into account, so it does not depend on which sequence was the query. Of the two alignments of a pair, the one with fewer differences was used in both directions; on a tie, the one with the higher bit score, and then the longer one, was kept. This makes the classification independent of the order of the search results: a substitution at the last position may be included in the alignment in one direction and left out in the other, and with the same number of differences the identity of the shorter alignment is higher (e.g. 98.1% instead of 97.9% with eight differences), which can move a pair across the 98% near-identical threshold. The effect of the thresholds was assessed in a sensitivity analysis: all pairs of distinct proteins (for each pair, the alignment direction with fewer differences) were reclassified with one threshold changed at a time, and we counted the proportion of pairs that moved to a different relation type. The difference count of the close pairs was decomposed into three components: amino acid substitutions within the alignment, internal gaps and unaligned ends.

**Table 3.** Relation types and the sequence identity and coverage thresholds that define them.

| Relation | Identity | Coverage |
|---|---|---|
| identical protein | 100% | 100% |
| near-identical | ≥98% | both ≥95% |
| length variant | ≥98% | one ≥95%, the other ≥80% |
| same family | ≥90% | both ≥80% |
| homologue | ≥50% | both ≥50% |

The mapping is directional, so the best hit was determined separately for every entry and target database. Can- didates were ranked first by relation type (identical protein, near-identical, length variant, same family, homologue), and within the same type by difference count. Since the best counterpart of A in database B need not have A as its own best counterpart, reciprocal best hits were flagged separately.

In allele-rich families, an entry often has several equally close counterparts in the target database. A new allele may, for example, differ by exactly one amino acid from two different alleles listed there. In such cases the sequence alone cannot determine which counterpart is the ‘correct’ one, so we did not choose one arbitrarily but retained all of them. A tie was declared if the candidates have the same relation type, exactly the same difference count, and are different proteins. Target entries encoding the same protein (e.g. synonymous nucleotide variants) counted as a single candidate. For every entry we recorded the number of equally good candidates and whether they belong to the same NCBI family (or, without a label, to the same sequence-based cluster). If they do, the mapping is ambiguous only at the allele level and unambiguous at the family level. If not, the uncertainty extends to the family level.

Close relations (identical protein, near-identical, length variant) were retained in full, because they provide the allele- and gene-level mapping. Of the weaker relations (same family, homologue), only the best 3 per entry and target database were kept. In large allele families (e.g. the *blaPDC* or *blaOXA* family), the number of weak relations grows quadratically with the number of entries, while beyond the closest ones they carry little information. Candidates tied at rank 3 (same relation type and difference count) were all retained, so the truncation does not depend on the order of the candidates. The list of weak relations is therefore incomplete. All hits of the search are available in the protein_hits.tsv file.

### 2.4 AMR gene families

By gene family (hereafter family) we mean a group of genes of common origin that are close to each other in sequence and typically share the same resistance mechanism, for example the TEM-type *β*-lactamases (*blaTEM*) or the *sul1* group of sulfonamide-resistant dihydropteroate synthases. The family is the middle level of organisation. Below it are the individual alleles, which often differ by only one or two amino acids (e.g. *blaTEM-1* and *blaTEM-116*), and above it the broader functional categories (e.g. class A *β*-lactamases). Family-level comparison matters because the allele-level mapping is often ambiguous or incomplete. A newly identified allele is usually not yet present in the other databases, but its family is.

The databases interpret the family at different levels. In the curated gene hierarchy of AMRFinderPlus, the family is the grouping level closest to the gene and the allele (e.g. *blaTEM* for allele-rich families, the gene itself for single genes, e.g. *dfrA1*). The *AMR Gene Family* categories of CARD are typically broader, mostly denoting an enzyme, transporter or mechanism family (e.g. *AAC(6’)*, *Erm-like 23S rRNA methyltransferase*, *major facilitator superfamily (MFS) antibiotic efflux pump*). ResFinder specifies no family, which can only be inferred from the gene names. Family membership was therefore determined in two complementary ways: by sequence-based family clusters and by family labels derived from the curated NCBI gene hierarchy.

#### Sequence-based family clusters

The unique proteins were sorted by decreasing length (at equal length by an identifier derived from the sequence digest, to make the result deterministic). Proceeding in this order, every protein not yet assigned became the representative of a new cluster, and its cluster received the unassigned proteins whose sequence identity *with the representative itself* is at least 90% and whose alignment covers at least 80% of the length of both proteins. A protein thus always joined the cluster of the first, that is the longest, representative to which it was close enough. The principle of the procedure is the same as that of the widely used CD-HIT [21] clustering method. Choosing the longest protein as representative prevents a shorter, truncated reference from becoming the representative, to which full-length proteins could not be linked because of the coverage requirement. The representative-based procedure was chosen over simple single-linkage clustering. With the latter, two proteins fall into the same cluster even if they are similar not to each other but only through an intermediate sequence, so related families can chain together, whereas in a representative-based cluster every member is directly similar to the representative. Cluster identifiers were derived from the sorted identifiers of the member proteins, so the identifier of a cluster does not change as long as its membership is unchanged.

#### NCBI family labels

The NCBI gene hierarchy is a tree-structured classification (ReferenceGeneHierarchy) whose *nodes* are nested, named gene groups. Below the root are the main categories (e.g. antimicrobial resistance, stress response), below these the groups by mechanism and function (e.g. the *β*-lactamases, within them the class A *β*-lactamases), then the families (e.g. *blaTEM*), and finally, as the leaves of the tree, the individual genes and alleles (e.g. *blaTEM-1*). Every node except the root has one parent, and the hierarchy also indicates whether a node is an allele. The Reference Gene Catalog of AMRFinderPlus gives for every entry the node of the hierarchy it belongs to (Fig. 2).

**Figure 2.**
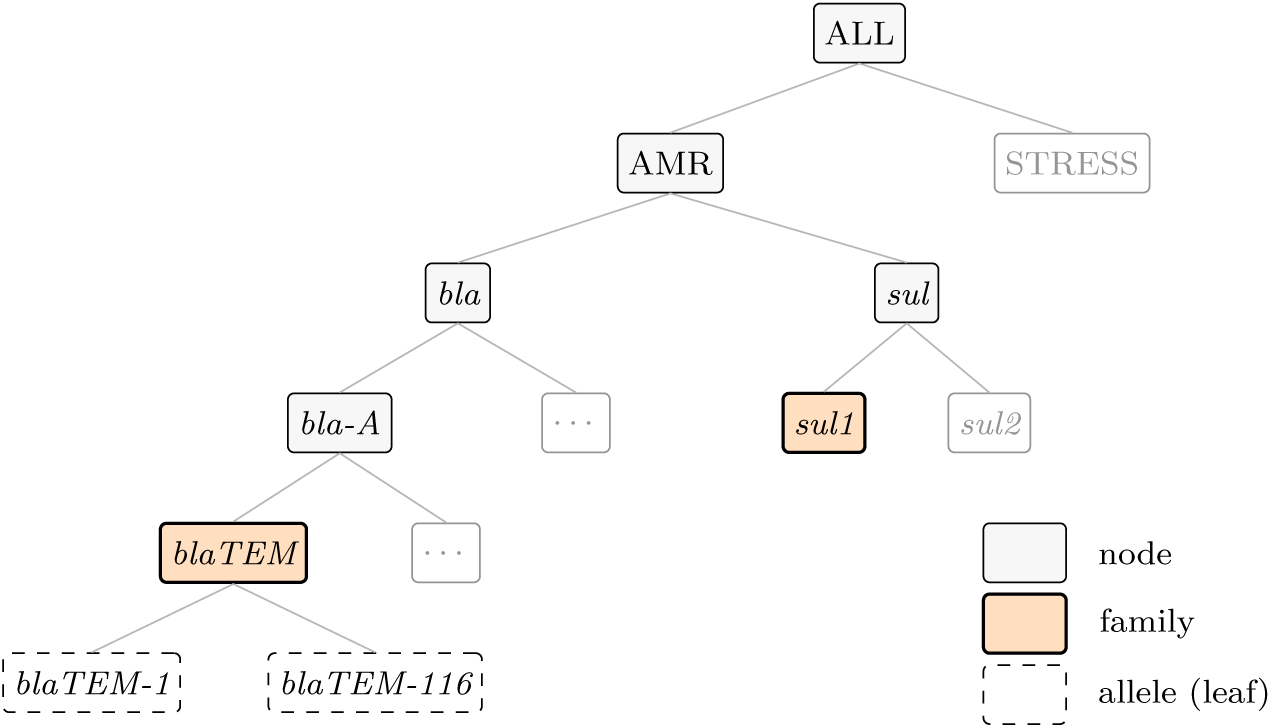
Part of the NCBI gene hierarchy. In allele-rich families the node of an entry is an allele (e.g. *blaTEM-1*), and the family is then the parent node (*blaTEM*); for single genes (e.g. *sul1*) the node of the entry is itself the family. Grey nodes indicate further branches of the hierarchy not detailed in the figure.

If the node is an allele (e.g. *blaTEM-1*), the family is the parent node (*blaTEM*); if it is not an allele but a gene-level node not subdivided into allele names (e.g. *sul1*), the node itself is the family. Family boundaries are set by the curators of the hierarchy, who take into account the evolutionary origin of the genes besides sequence similarity. The entries of CARD, ResFinder and DisinFinder have no NCBI node (CARD’s own family categories, as seen above, group at a different level), so their family labels were derived from the AMRFinderPlus entries, which allows the same family concept to be used for every database. If the protein is also encoded by an AMRFinderPlus entry, its family was adopted. If not, we considered the AMRFinderPlus proteins that share at least 90% identity with it, with an alignment covering at least 80% of the length of both proteins. Of these, the family of the closest one, that is the one differing in the fewest amino acids, was used. If the identity with the protein of the closest and of the second closest family differed by less than 1 percentage point, the derived label was flagged as ambiguous, because the family assignment is then not unambiguous from the sequence. If there was no such AMRFinderPlus protein, the protein received no family label.

#### Comparison of the family concepts

On the AMRFinderPlus entries, we compared the sequence-based clusters with the NCBI families. An NCBI family was considered *split* if its members fell into more than one cluster, and a cluster *merged* if it contained members of more than one NCBI family. This shows where the uniform identity threshold departs from the curated family boundaries. The agreement was also quantified over the threshold range of 85–98% with the adjusted Rand index [22], repeating the clustering at every threshold. To characterise the relationship between the CARD families and the NCBI hierarchy, for every CARD *AMR Gene Family* category we collected the NCBI family labels of its entries (directly if the protein is also encoded by an AMRFinderPlus entry, otherwise derived as described above). We then determined the lowest common ancestor of these families in the NCBI hierarchy, that is, the deepest node that is an ancestor of all of them, and how many levels above the families it lies on average. The topmost node of the hierarchy and the main antimicrobial resistance branch were considered the root, because a common ancestor below them does not yet represent an actual grouping. Since the strict common ancestor may fall to the root because of a few members classified elsewhere, a *majority common ancestor* was also determined, that is, the deepest node below which at least 80% of the NCBI families of the CARD family fall. If the strict common ancestor is the root but the majority one is not, the discrepancy is caused by a few outlying members. If, however, the majority common ancestor is also the root, CARD places in one family genes that the NCBI hierarchy does not group under a common node.

### 2.5 Drug class harmonisation

The databases use different class names. AMRFinderPlus, for example, gives broad classes (BETA-LACTAM), which CARD subdivides further (penicillin beta-lactam, cephalosporin, carbapenem). For AMRFinder- Plus, the class names were taken from the class field of the Reference Gene Catalog, where combined values (e.g. LINCOSAMIDE/STREPTOGRAMIN) denote several classes. For CARD, the ontology drug class categories assigned to the models were used, and for ResFinder and DisinFinder the class column of the phenotype table or, if missing, the name of the source file (e.g. beta-lactam.fsa). An entry may thus belong to several classes. These class names were assigned to 37 common classes (118 terms) with a manually editable mapping table published with the tool. Some examples from the table are shown in Table 4.

**Table 4.** Selected examples of the mapping of native drug class terms to the common classes, with the terms that occur in the presented release, ordered by frequency (at most four per cell). An asterisk marks ResFinder entries classified by their file name.

| Common class | AMRFinderPlus | CARD | ResFinder, DisinFinder |
|---|---|---|---|
| Beta-lactam | BETA-LACTAM | cephalosporin, penicillin beta-lactam, carbapenem, monobactam | Beta-lactam |
| Polymyxin/peptide | COLISTIN, LIPOPEPTIDE, BACITRACIN | peptide antibiotic, nonribosomal peptide antibiotics | Polymyxin |
| Sulfonamide | SULFONAMIDE | sulfonamide antibiotic | sulphonamide* |

Harmonisation necessarily merges finer subdivisions; the *β*-lactam subclasses of CARD, for example, were placed in a single *β*-lactam class. For ambiguous terms, the name of the source file decided. The ResFinder class ‘Folate pathway antagonist’, for example, may denote either sulfonamide or trimethoprim resistance, so for these entries the sulfonamide or trimethoprim source file determined the class. Terms that do not denote a class were ignored. Examples are the CARD labels *azole antibiotic* and *imidazole antibiotic*. In the database version used, they are assigned exclusively to entries of CTX-M *β*-lactamases, always alongside the cephalosporin class, so they do not denote a separate resistance class. As the classification of the databases may change with updates, the content report of each ARGeement release lists the new class names not yet present in the mapping table, and shows which gene families the ignored terms belong to, so that it becomes visible if an update also assigns them to other genes.

### 2.6 Name concordance

The relationship between names and sequences was examined from two directions. In the first approach, we took the cross-database entry pairs that encode the *same protein* and examined how well their names agree. Every pair was assigned to the first matching category of Table 5, that is, a pair was placed in a later category only if none of the earlier ones explained the difference. The first six categories arise from known naming conventions and can be handled by rules. The last two are genuine naming conflicts, in which the same protein is denoted by incompatible names.

**Table 5.**
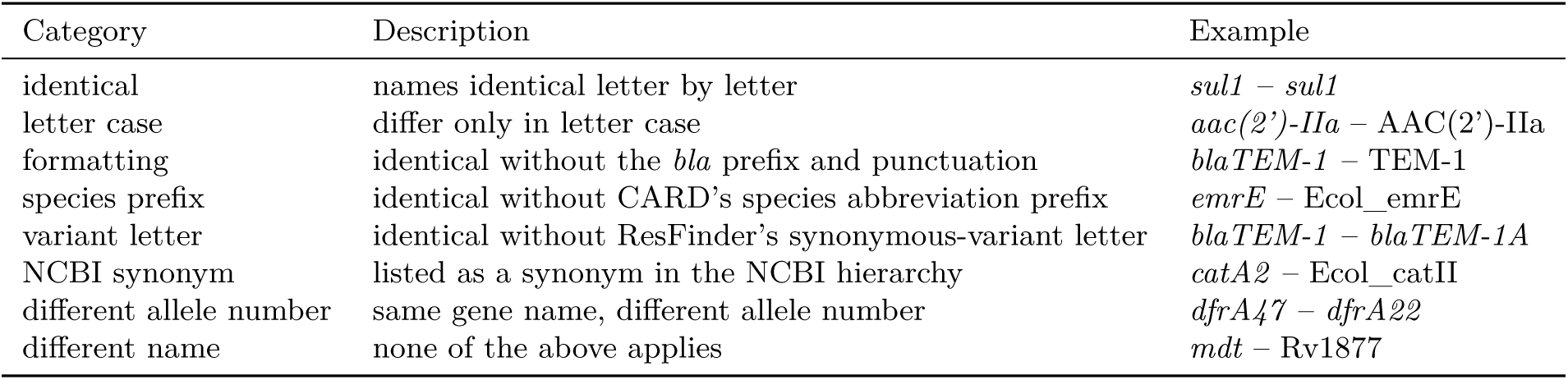
Name concordance categories for entry pairs encoding the same protein. The last two categories are genuine naming conflicts.

| Category | Description | Example |
|---|---|---|
| identical | names identical letter by letter | *sul1* – *sul1* |
| letter case | differ only in letter case | *aac(2')-IIa* – AAC(2')-IIa |
| formatting | identical without the *bla* prefix and punctuation | *blaTEM-1* – TEM-1 |
| species prefix | identical without CARD's species abbreviation prefix | *emrE* – Ecol_emrE |
| variant letter | identical without ResFinder's synonymous-variant letter | *blaTEM-1* – *blaTEM-1A* |
| NCBI synonym | listed as a synonym in the NCBI hierarchy | *catA2* – Ecol_catII |
| different allele number | same gene name, different allele number | *dfrA47* – *dfrA22* |
| different name | none of the above applies | *mdt* – Rv1877 |

In the second approach, we took the cross-database pairs whose names are equivalent under the rules of the first five categories, and examined how well their proteins agree. NCBI synonyms were not used in this direction, because we examine the behaviour of identical names, whereas synonyms link different names; moreover, they belong only to AMRFinderPlus entries, so they cannot be applied between the names of CARD and ResFinder. Their relation was assigned to the categories of Table 3 based on all hits of the all-against-all comparison. They were considered *unrelated* if there was no hit between them, that is, if their sequence identity remained below 50% or the alignment covered less than 50% of the length of either protein. Based on the NCBI hierarchy, we also recorded whether the name on the AMRFinderPlus side denotes an allele or a gene-level group. For a gene-level name, under which AMRFinderPlus lists several sequence variants without separate allele names (e.g. *sul1*), a different sequence is expected, whereas for an allele name it indicates a genuine conflict. ORF sub-entries obtained by splitting were left out of both approaches, because they carry the name of the operon.

### 2.7 Mutation inventory

The wild-type and mutant sequences of mutation-based determinants are almost identical, so they cannot be compared by sequence identity, and they were left out of the mapping. Instead, they were inventoried by genus in all three sources. In the protein mutation table and the per-taxon-group rRNA mutation tables of AMRFinderPlus, every row describes a mutation; these were counted by the genus of the taxon group and the mutated gene. In CARD, mutations are described by variant models (protein and rRNA variant models); for these the species of the model’s reference sequence gave the genus, and the point mutations listed in the models were counted. In the per-species directories of PointFinder, the genes carrying mutations and the mutation positions were counted. The units differ between sources (mutation, point mutation listed in a model, and mutation position, respectively), so the numbers primarily show which genera the mutation part of each database covers, and are only roughly comparable with each other.

### 2.8 Synthetic validation

The accuracy of the mapping was also checked on synthetic variants created with known modifications (argeement validate --n 500 --seed 1). From the curated (AMRFinderPlus and CARD) proteins of the release, we randomly selected 500 proteins of 100–1,000 amino acids starting with methionine. From each, four protein-level variants were created: an exact copy, a variant with 1–5 random amino acid substitutions, a variant extended N-terminally by 3–10 amino acids, and one truncated C-terminally by 3–10 amino acids. Nucleotide-level variants were also created. A CDS encoding the protein, assembled from randomly chosen synonymous codons, was used on the forward strand, on the reverse strand and without its start codon, and two composite sequences were also built. One is a two-gene operon with the next protein of the sample; in the other, the gene is accompanied by an unrelated 90-amino-acid ORF with no similarity to resistance genes. The variants were placed in the release with the same procedure (translation, ORF splitting, MMseqs2 search, classification) as the database entries. Four metrics were computed for each variant type. Recovery shows whether the original protein is among the best hits (including ties), and uniqueness whether it is the single best hit. The relation type and the difference count are correct if they equal the value expected from the modification under the rules of Table 3.

### 2.9 Real-world use case

The practical value of the mapping was examined on real genomes: on 25 randomly selected complete RefSeq genomes from each of four species (*Escherichia coli*, *Klebsiella pneumoniae*, *Salmonella enterica*, *Staphylococcus aureus*), drawn with a fixed random seed (concordance/select_genomes.py --n 25 --seed 1). Because the selection depends on the RefSeq genomes available at the time, the list of the selected genomes (concordance/genomes.tsv in the GitHub repository) is the reference for reproduction. The genomes were annotated with AMRFinderPlus (4.2.7), the Resistance Gene Identifier (RGI; 6.0.8; CARD) and ResFinder (4.7.2; together with DisinFinder), with default settings and exactly the database versions from which the release described here was built. All three tools aligned with BLAST (2.16.0): ResFinder uses it by default on assembled genomes (KMA is intended for raw reads), and for RGI we selected it instead of the default DIAMOND. RGI was run in protein mode on ORFs predicted with Prodigal (2.6.3) [23]; contig mode works the same way but does not handle RefSeq contig names correctly. The comparison included calls of the same scope as the mapping: the acquired resistance and biocide resistance genes of AMRFinderPlus (without point mutations), the protein homolog models of RGI, and all hits of ResFinder and DisinFinder. Two calls were considered the same locus if they lie on the same contig and their overlap is at least 80% of the length of both calls (reciprocal overlap); equally good references reported by ResFinder at the same position form a single locus. For loci called by only one tool of a pair, we used the release to check whether the reported reference has a close counterpart (identical protein, near-identical or length variant) in the database of the other tool, to separate differences in database content from differences in detection. The locus-level results are available in the GitHub repository (concordance/loci_2026-11-16_184628.tsv).

The calls of each tool pair were compared in three ways. *Locus-level agreement* measures whether the two tools find a resistance gene at the same place: the share of loci called by both among the loci called by at least one of them. At the loci called by both tools, we also examined whether the two tools name the same gene. For *agreement before mapping*, only the gene names were compared: a locus was considered concordant if the two names are identical ignoring letter case. For *agreement after mapping*, the relation between the reference entries given by the two tools was taken from the release: a locus was considered concordant if the two references are the identical protein, near-identical or length variants. Loci called by both tools that remained discordant after mapping were classified by the relation of the two references: the two references belong to the same family, or there is no close relation between them. A separate category would have held loci at which one of the references cannot be linked to any usable protein of the release (e.g. because of a different database version or an entry excluded from the mapping); no such locus occurred in the genomes examined.

## 3 Results

### 3.1 Content of the databases

Of the 17,920 entries of the release described here (2026-11-16), 39 could not be translated as a single coding sequence, because they contained several genes or were stored on the reverse strand or in a shifted frame. These were split into ORFs, so that a total of 17,959 protein sequences (protein-level entries) entered the comparison, encoding 9,337 unique proteins (Fig. 3). For 3 of the split entries no ORF was retained; these entries take no part in the protein-level comparison (Section 3.4). AMRFinderPlus is the largest (8,633 entries); CARD contains 6,059, ResFinder 3,212 and DisinFinder 16 entries.

**Figure 3.**
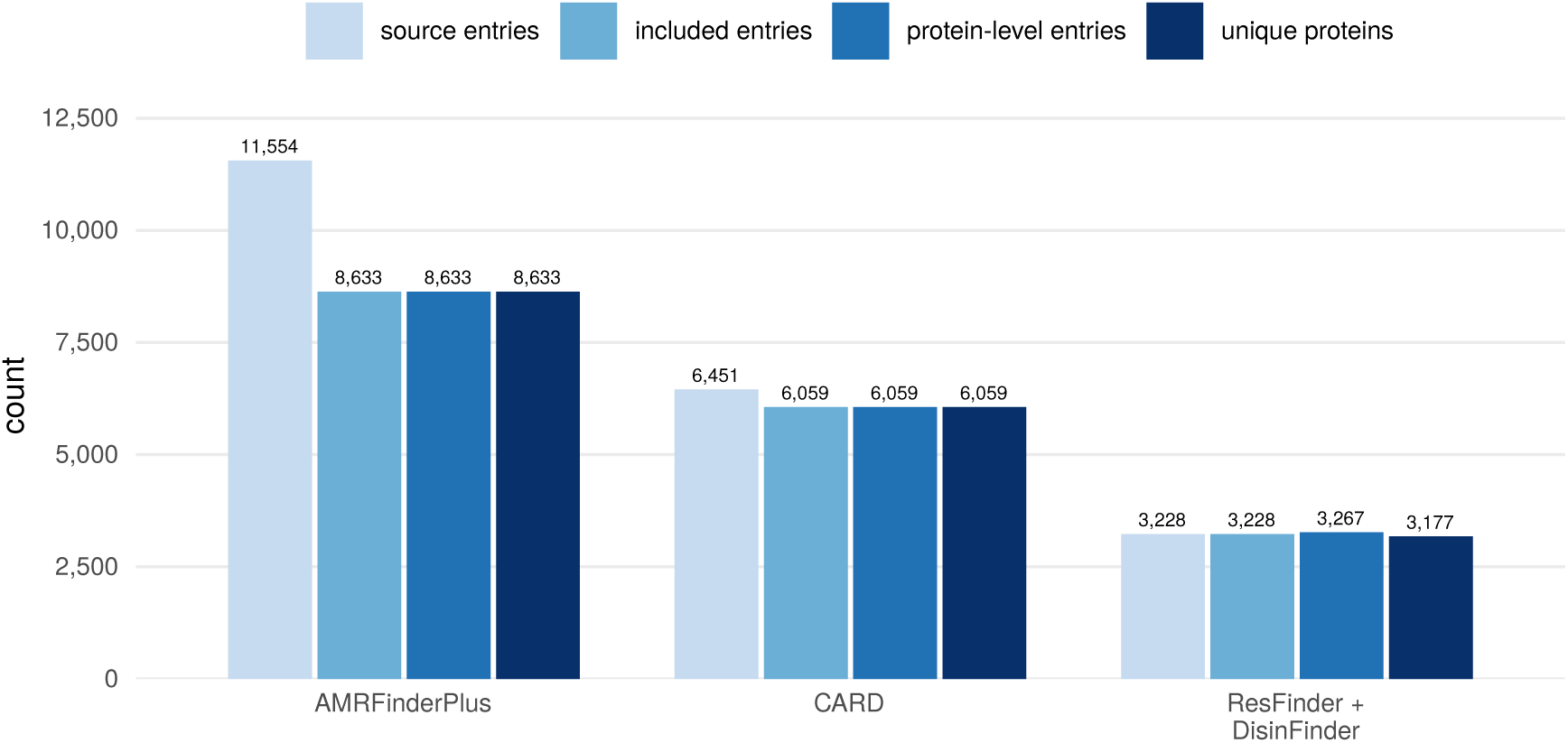
Size of the databases. Number of entries in the source, of entries included in the mapping, of protein-level entries and of unique proteins. ResFinder and DisinFinder (Center for Genomic Epidemiology, CGE) are shown together; as some proteins are encoded by both databases, the number of unique CGE proteins is slightly smaller than the sum for the two databases. The source count of AMRFinderPlus includes all rows of the Reference Gene Catalog, together with the rows of mutation, virulence and stress response genes.

The distribution by harmonised drug class is shown in Fig. 4. *β*-Lactams dominate in all three databases (AMRFinderPlus 6,228, CARD 4,980, ResFinder 2,030 entries), a consequence of the large allele families (e.g. TEM, CTX-M, OXA). In the other classes the databases show different emphases. AMRFinderPlus lists considerably more aminoglycoside resistance genes (719) than CARD (239), and AMRFinderPlus also leads for glycopeptides (236 and 82, respectively). CARD has more entries assigned to polymyxin and peptide resistance (234 and 135, respectively) and to rifamycin resistance (38 and 21), whereas for sulfonamides ResFinder is the largest (54 entries, compared with 41 in AMRFinderPlus and 10 in CARD). A few rare classes occur in only one database, for example aminocoumarins only in CARD and bleomycin only in AMRFinderPlus. Some of these are genuine differences in content, others arise from the classification approach. AMRFinderPlus labels some efflux pumps with a general, multidrug class, which has no counterpart in CARD, where efflux pumps are assigned specific drug classes. Biocide resistance genes are present in CARD (62), AMRFinderPlus (40) and DisinFinder (17), but, in line with the division of work within CGE, not in ResFinder.

**Figure 4.**
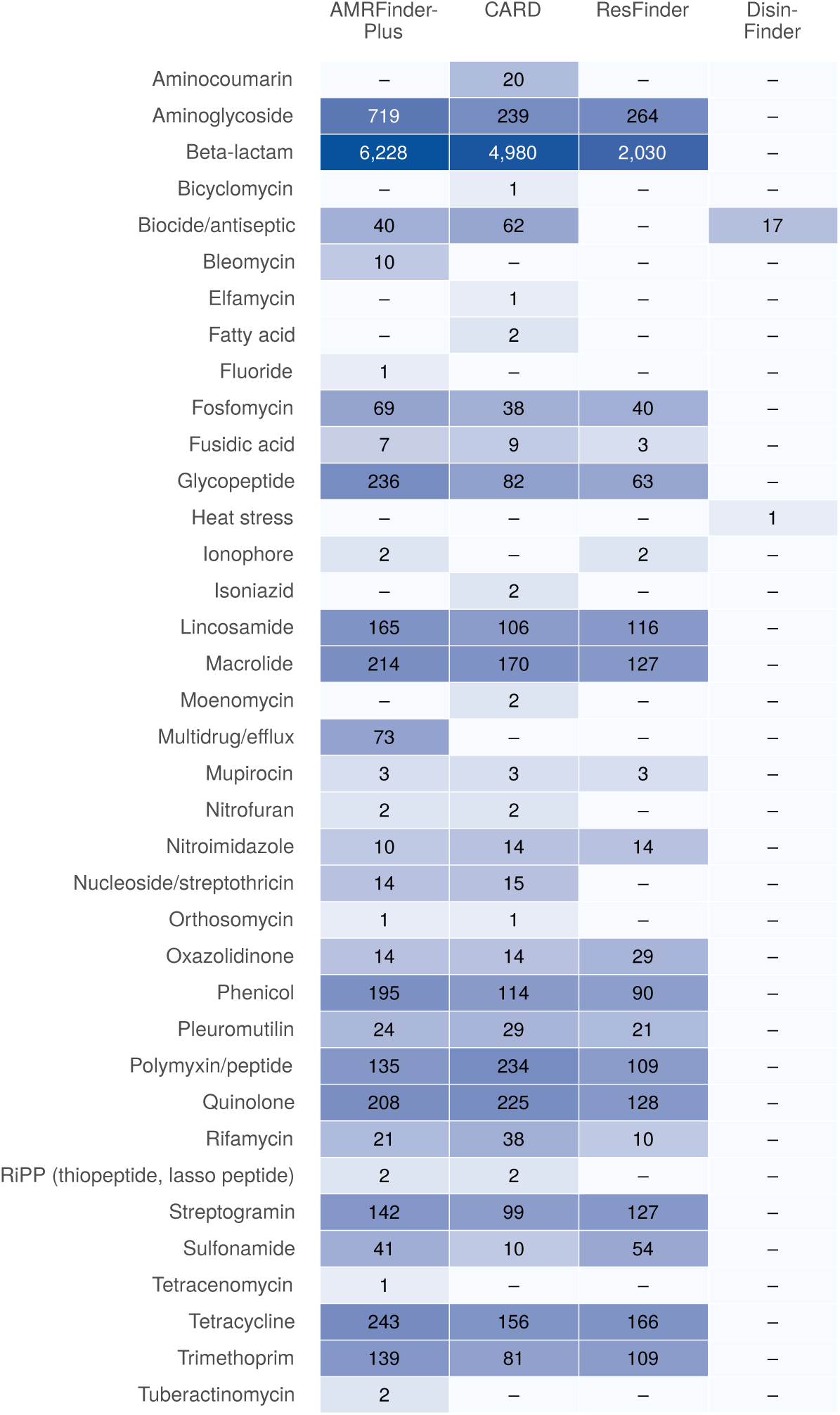
Number of protein-level entries by harmonised drug class. An entry appears in every class assigned to it. The figure shows all 37 harmonised classes, in alphabetical order.

### 3.2 Overlap between the databases

At the protein level, 6,168 (66.1%) of the unique proteins occur in at least two databases, and 2,364 (25.3%) in all three (counting ResFinder and DisinFinder together). At the entry level, the overlap is direction-dependent, because the shares always refer to the entries of the source database, and the mapping is not one-to-one (Fig. 5). In AMRFinderPlus, 96.5% of the ResFinder entries and 96.1% of the CARD entries have a close counterpart, and 89.6% and 93.4%, respectively, an identical one. Conversely, only 88.8% of the AMRFinderPlus entries are closely linked to CARD and 77.6% to ResFinder. AMRFinderPlus thus covers most of the included content of the other two databases, while also containing many alleles that are absent from them. These shares, however, reflect not only differences in content but also the different inclusion scopes of the databases (Table 2). ResFinder lists only acquired resistance genes, whereas the protein homolog models of CARD include many chromosomal genes conferring intrinsic resistance, for example components of efflux pumps. In AMRFinderPlus these belong partly to the plus set or to the stress response genes, and are partly missing. Directional coverage can therefore only be interpreted under the given inclusion rules. After stratification (Table 6), 99.6% of the antibiotic-inactivating entries of CARD have a close counterpart in AMRFinderPlus, but only 35.9% of the 276 entries with an efflux mechanism, and none of the 12 entries with reduced permeability. Most of the CARD entries without a close AMRFinderPlus counterpart are thus efflux and permeability genes, which the two databases handle differently. Of the core entries of AMRFinderPlus, 89.5% are closely linked to CARD and 78.7% to ResFinder; of the plus entries, 38.1% and 5.6%, respectively. Plus entries are thus much less often present in the other two databases, but they make up only a small part of the content of AMRFinderPlus (Table 6). No other database has a close counterpart for 747 (8.7%) of the AMRFinderPlus entries, 224 (3.7%) of the CARD, 100 (3.1%) of the ResFinder and 7 (38.9%) of the DisinFinder entries.

**Figure 5.**
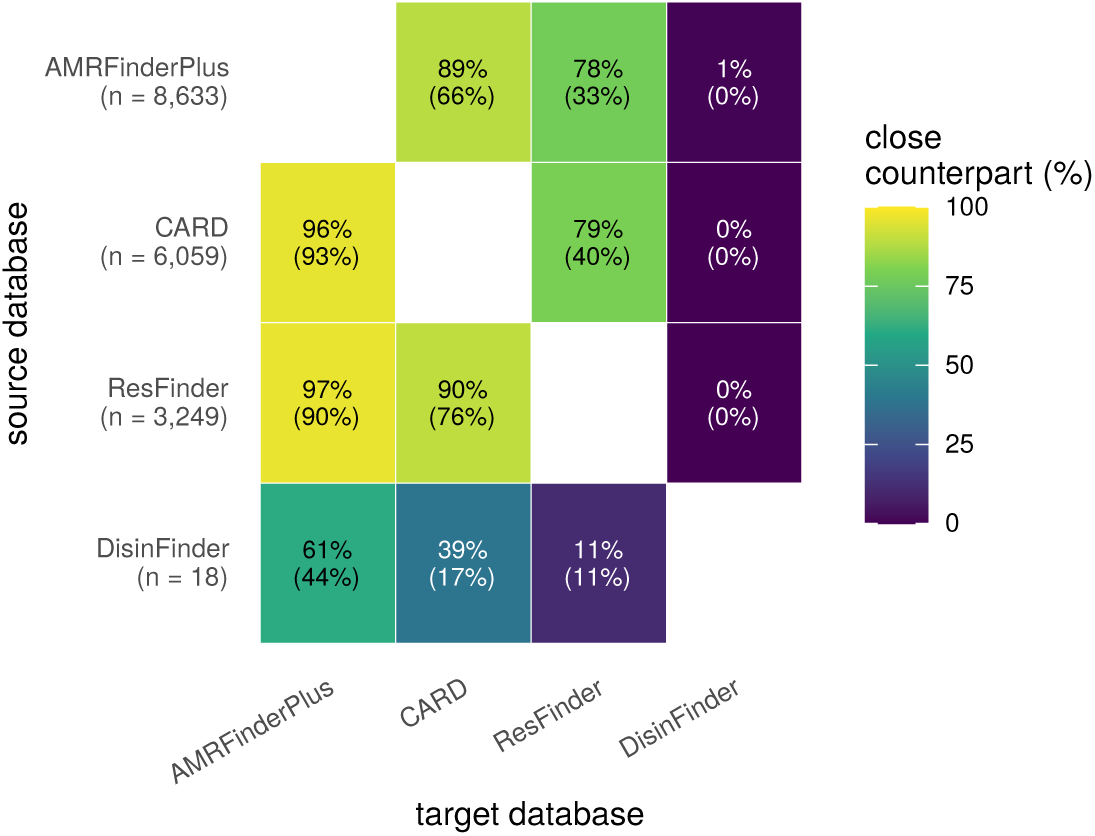
Overlap between the databases. Each cell shows the percentage of the entries of the database in the row that have a close counterpart (identical protein, near-identical or length variant) in the database in the column. The value in parentheses counts only counterparts encoding the identical protein. For example, the AMRFinderPlus column of the ResFinder row shows the share of ResFinder entries that have a close or identical-protein counterpart in AMRFinderPlus. The figure is not symmetrical, because the shares always refer to the entries of the database in the row, and several entries of one database may correspond to the same entry in the other.

**Table 6.** Stratified directional coverage. The share of CARD entries with a close counterpart in AMRFinderPlus by resistance mechanism (an entry may belong to more than one mechanism), and the share of AMRFinderPlus entries with a close counterpart in CARD and ResFinder by whether AMRFinderPlus places the entry in its core or plus set. The set was taken from all catalogue rows of the protein accession: an entry belongs to the core set if any of its rows is core.

| Direction | Stratum | *n* | Close counterpart |
|---|---|---|---|
| CARD → AMRFinderPlus | antibiotic inactivation | 5,245 | 99.6% |
| CARD → AMRFinderPlus | antibiotic target alteration | 293 | 90.1% |
| CARD → AMRFinderPlus | antibiotic efflux | 276 | 35.9% |
| CARD → AMRFinderPlus | antibiotic target protection | 170 | 98.2% |
| CARD → AMRFinderPlus | antibiotic target replacement | 72 | 95.8% |
| CARD → AMRFinderPlus | reduced permeability to antibiotic | 12 | 0.0% |
| CARD → AMRFinderPlus | resistance by host-dependent nutrient acquisition | 1 | 100.0% |
| AMRFinderPlus → CARD | core set (core) | 8,507 | 89.5% |
| AMRFinderPlus → ResFinder | core set (core) | 8,507 | 78.7% |
| AMRFinderPlus → CARD | plus set (plus) | 126 | 38.1% |
| AMRFinderPlus → ResFinder | plus set (plus) | 126 | 5.6% |

### 3.3 Names and sequences

For 64.4% of the 10,954 cross-database entry pairs encoding the same protein, the names differ even when letter case is ignored, so a naive name-based comparison would not recognise these pairs (Fig. 6a). Most differences arise from naming conventions. These include formatting differences (6,526 pairs, mainly the *β*-lactamase names of CARD without the *bla* prefix), the species prefixes of CARD (20), the variant letters of ResFinder (34) and NCBI synonyms (32). A genuine naming conflict was found for 447 pairs. Of these, in 66 cases a different allele number and in 381 cases an entirely different name denotes the same protein.

**Figure 6.**
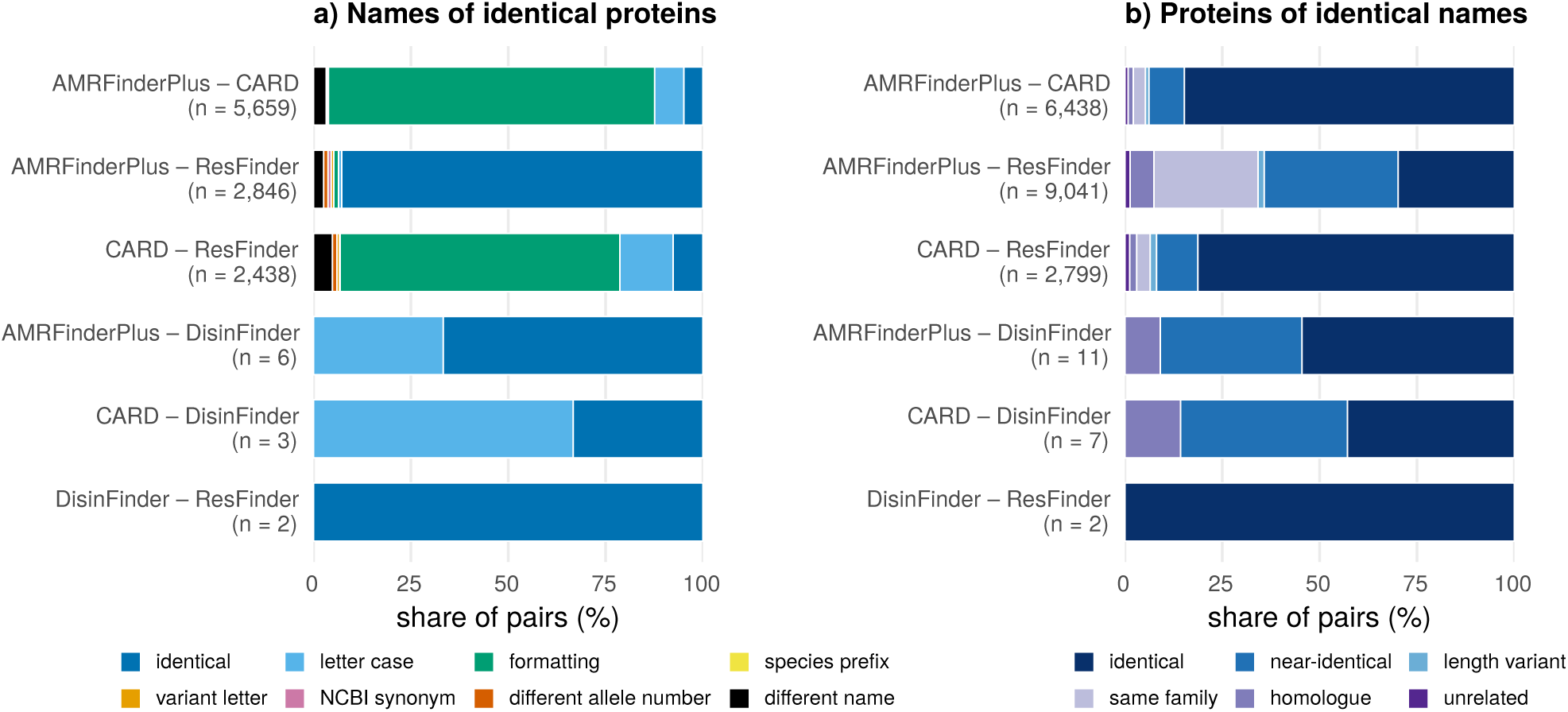
Name concordance. a) Names of identical proteins. Names of cross-database entry pairs encoding the same protein, by category. b) Proteins of identical names. Relation of the proteins of cross-database entry pairs with the same name.

Of the 18,298 cross-database pairs with the same name, 42.9% do not encode the same protein (Fig. 6b), 693 pairs are only distant homologues, and 209 pairs are not even related by sequence. Of the 7,129 pairs named with an allele name, 146 (2.0%) do not encode the same protein, and of these 38 are linked at most at the same-family level.

### 3.4 Operons and shifted reading frames

In total, 34 ResFinder and 5 DisinFinder entries had to be split into ORFs. Splitting yielded 74 ORFs from these in ResFinder and 15 in DisinFinder. Of these, 71 and 7, respectively, were retained in the mapping, and 3 and 8 were dropped, because they were not linked to a curated protein and the splitting pattern did not justify retaining them either. Most split entries are operons containing several genes; the others are genes stored on the reverse strand (e.g. *aac(3)-I*, *aac(3)-Xa*, *blaBKC-1*, *mph(A)*, *qepA1*, *grdA*, *blaSPG-1*), which splitting recovered as a single full-length ORF. Each of the six *vanHAX* operon entries was split into its three genes, and VanH (322 amino acids), VanA (343 amino acids) and VanX (194–205 amino acids) were retained in all six operons. Of the 18 genes, 17 gave a protein identical to an AMRFinderPlus entry, whereas in CARD only the genes of two operons have an identical counterpart (in one of them, except for VanA). The excluded ORFs fall into two groups. One consists of unrelated ORFs selected alongside the true gene, which come from neighbouring or opposite-strand stretches of the reference sequence (e.g. in the *qacC*, *qacD* and *qacH* entries). The other comprises the 3 entries (*mph(D)*, *blaFLC-1*, *qacF*) none of whose possible ORFs of at least 60 amino acids is linked to a curated protein, so they take no part in the mapping at the protein level. The reference sequence of *mph(D)* contains an internal stop codon, and the length of *blaFLC-1* is not even divisible by three, so in these the gene probably cannot be read in full length because of a frameshift. In this release, the curated-priority reselection did not change the ORFs of any entry, that is, in no case did a longer unrelated ORF displace a gene linked to a curated protein.

### 3.5 Families

Of the 1,477 sequence-based family clusters obtained on the 8,620 NCBI-labelled proteins of AMRFinderPlus, 83 span more than one NCBI family, and of the 1,355 NCBI families in total, 130 are split across more than one cluster. On all 9,337 unique proteins, 1,747 clusters were formed. For the chromosomal *β*-lactamases of *Klebsiella* species (SHV,

LEN and OKP), the NCBI hierarchy distinguishes 6 families (SHV 216, LEN 200, OKP-B 196, OKP-A 116, OKP-D 8, OKP-C 1 proteins). The lowest identity between protein pairs within a family is 91.6% (5th percentile 96.5%), whereas the highest identity between pairs from different families is 96.8%, and 28.1% of the between-family pairs reach the 90% same-family threshold (Fig. 7). Representative-based clustering assigned the 6 families to 4 clusters, and none of the clusters coincided with an NCBI family. All OKP-A proteins and most OKP-B proteins merged into one cluster (C2); the whole LEN family and most SHV proteins fell into a common cluster (C1), together with a few OKP-B, OKP-C and OKP-D proteins; the remaining SHV proteins formed a separate, smaller cluster (C3), and the fourth cluster (C4) consisted of two OKP-D proteins (Table 7). The agreement between the clusters and the NCBI families on this subset is moderate (adjusted Rand index 0.440). The NCBI family label was available directly for 8,620 proteins and derived for 435 proteins; of the latter, 28 labels are ambiguous, because the identities of the two closest families differ by less than 1 percentage point.

**Figure 7.**
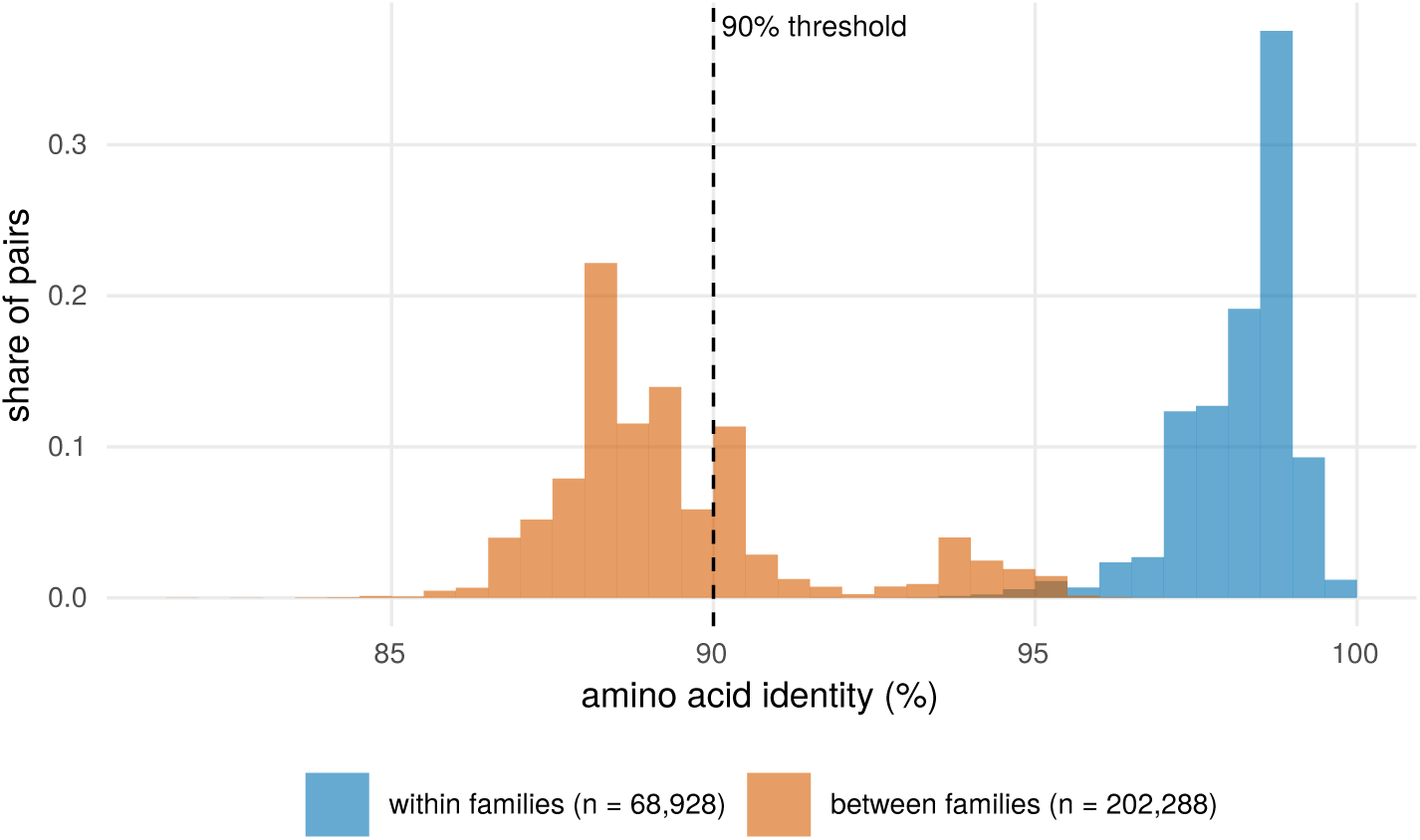
Amino acid identity of all protein pairs of the 6 NCBI families of the chromosomal *β*-lactamases of *Klebsiella* species (SHV 216, LEN 200, OKP-B 196, OKP-A 116, OKP-D 8, OKP-C 1 proteins), within families (68,928 pairs) and between families (202,288 pairs), as relative frequencies. The dashed line is the 90% identity threshold of the same family; classification also requires a coverage of at least 80% on both proteins.

**Table 7.** Chromosomal *β*-lactamases of *Klebsiella* species: distribution of the AMRFinderPlus proteins by NCBI family (rows) and sequence-based family cluster (columns). Clusters (C1–C4) are numbered in decreasing order of size. If the two partitions agreed, every row and every column would contain a single filled cell. The adjusted Rand index between the two partitions is 0.440.

| NCBI family | C1 | C2 | C3 | C4 | Total |
|---|---|---|---|---|---|
| *blaLEN* | 200 | – | – | – | 200 |
| *blaOKP-A* | – | 116 | – | – | 116 |
| *blaOKP-B* | 12 | 184 | – | – | 196 |
| *blaOKP-C* | 1 | – | – | – | 1 |
| *blaOKP-D* | 6 | – | – | 2 | 8 |
| *blaSHV* | 190 | – | 26 | – | 216 |
| Total | 409 | 300 | 26 | 2 | 737 |

Of the 388 CARD *AMR Gene Family* categories, 366 had an NCBI family label; 289 of these (79.0%; 74.5% of all categories) corresponded to exactly one NCBI family (Table 8). For most CARD families spanning more than one NCBI family, the common ancestor is an intermediate node of the NCBI hierarchy (e.g. dfr_gen for the *trimethoprim resistant dihydrofolate reductase dfr* family). In 7 cases, the strict common ancestor is the root of the hierarchy, but the majority common ancestor is an intermediate node. Of the NCBI families in the *Erm-like 23S rRNA methyltransferase* family, 93% lie below a common node, and of those in the *major facilitator superfamily*

**Table 8.**
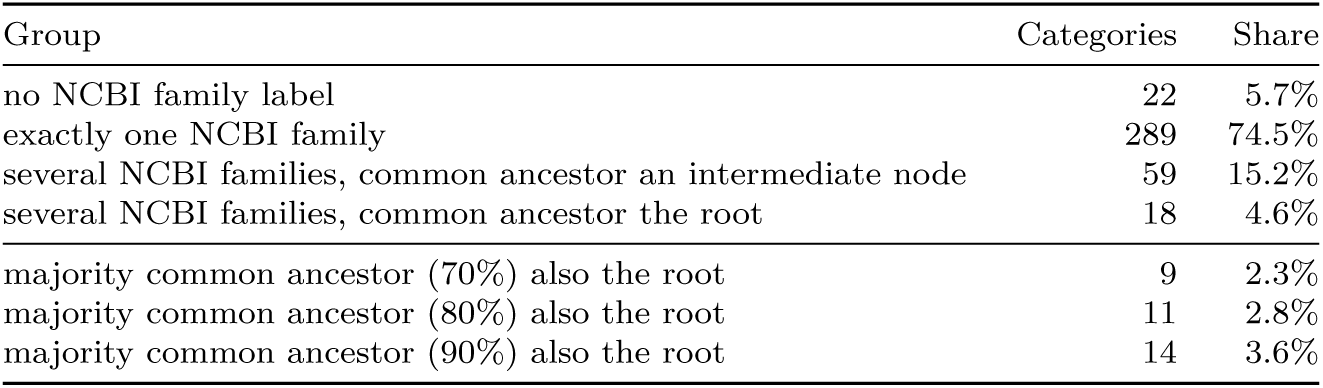
Relationship of the 388 CARD *AMR Gene Family* categories to the NCBI hierarchy. The upper four rows partition the categories by the NCBI family labels of their entries and the strict common ancestor of these families. The lower rows show for how many categories the majority common ancestor also remains at the root when 70, 80 or 90% of the families must fall below it.

| Group | Categories | Share |
|---|---|---|
| no NCBI family label | 22 | 5.7% |
| exactly one NCBI family | 289 | 74.5% |
| several NCBI families, common ancestor an intermediate node | 59 | 15.2% |
| several NCBI families, common ancestor the root | 18 | 4.6% |
| majority common ancestor (70%) also the root | 9 | 2.3% |
| majority common ancestor (80%) also the root | 11 | 2.8% |
| majority common ancestor (90%) also the root | 14 | 3.6% |

*(MFS) antibiotic efflux pump* family, 87%. In 11 cases, the majority common ancestor is also the root; examples are the *Van ligase* and *glycopeptide resistance gene cluster* families, the nitroimidazole reductases and the ABC-type efflux pumps. The complete partition of the CARD categories, including the 22 categories without an NCBI family label, is shown in Table 8. The number of majority common ancestors remaining at the root is 9 at a majority threshold of 70% and 14 at 90%.

The adjusted Rand index measuring the agreement between the sequence-based clusters and the NCBI families was 0.886 at the default setting (90%) and varied between 0.645 and 0.915 over the threshold range of 85–98% (Table 9). The highest agreement was obtained at 92% (0.915), so the default setting is close to the threshold that best fits the curated family boundaries. With stricter settings more and more NCBI families are split, and with looser ones more and more clusters merge different families.

**Table 9.** Agreement between the sequence-based family clusters and the NCBI families at different identity thresholds, on the 8,620 unique AMRFinderPlus proteins (1,355 NCBI families). The coverage requirement was kept at 80% on both proteins. An NCBI family is split if its members fall into more than one cluster; a cluster is merged if it contains members of more than one NCBI family. ARI: adjusted Rand index (1 = perfect agreement, 0 = random).

| Threshold | Clusters | Split families | Merged clusters | ARI |
|---|---|---|---|---|
| 85% | 1,320 | 83 | 110 | 0.826 |
| 88% | 1,401 | 113 | 101 | 0.845 |
| 90% | 1,477 | 130 | 83 | 0.886 |
| 92% | 1,561 | 155 | 73 | 0.915 |
| 95% | 1,777 | 219 | 57 | 0.883 |
| 98% | 2,560 | 340 | 41 | 0.645 |

### 3.6 Mutation-based determinants

The mutation inventory contains data for 49 genera (Fig. 8). AMRFinderPlus lists 1,637 mutations in 20 genera, the variant models of CARD list 2,117 point mutations in 43 genera, and PointFinder 1,346 mutation positions in 10 genera. All three sources contain mutation data for 8 genera (*Campylobacter*, *Enterococcus*, *Escherichia*, *Helicobacter*, *Klebsiella*, *Neisseria*, *Salmonella* and *Staphylococcus*), whereas 33 genera occur in only one source (5 only in AMRFinderPlus, 27 only in CARD, 1 only in PointFinder). The genus occurring only in PointFinder is *Plasmodium*, because, besides bacteria, PointFinder also lists mutations conferring resistance to antimalarial drugs. CARD likewise includes two non-bacterial genera (*Chlamydomonas* and *Halobacterium*; Fig. 8).

**Figure 8.**
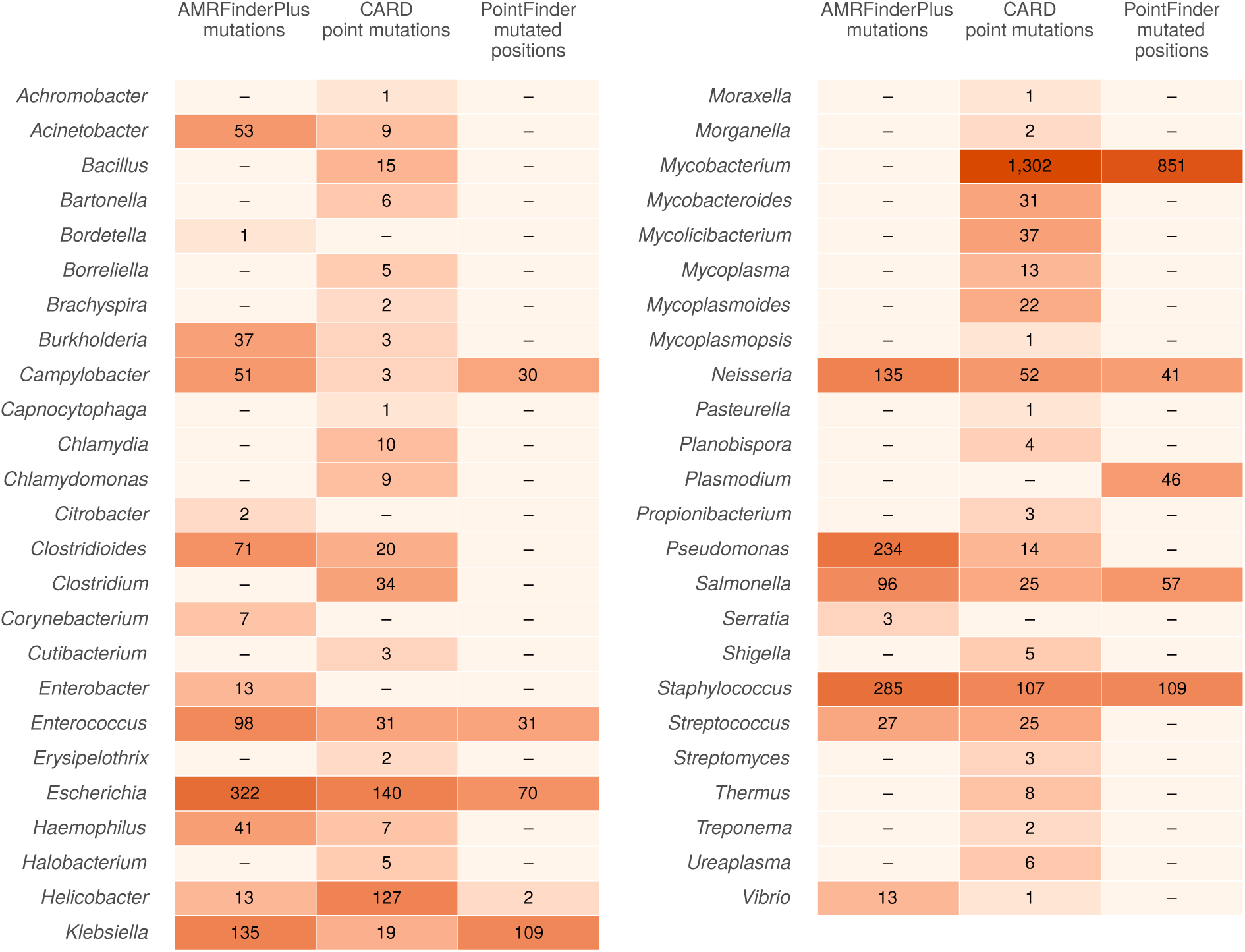
Mutation-based determinants by genus in the three sources. The AMRFinderPlus column counts the listed mutations, the CARD column the point mutations listed in the variant models, and the PointFinder column the mutation positions. As PointFinder lists several resistant amino acids at one position as a single item, the numbers are only roughly comparable between the columns, but all three show the genus coverage exactly.

### 3.7 Changes between two database versions

Between the release of the pinned profile amrfinder-2026-08-07 (AMRFinderPlus 2026-08-07.1) and the latest release (AMRFinderPlus 2026-10-29.1), 80 new entries appeared and 9 entries were renamed; no entry was removed, and the sequence of no entry changed under the same identifier. The release of the pinned profile (2026-11-16_203603) is deposited in a separate Zenodo record (10.5281/zenodo.24301243). Of the new entries, 35 have no close counterpart in any other database. The new and renamed entries all came from AMRFinderPlus; ResFinder also changed (2.6.0 → 2.7.0), but without adding or renaming entries. The relations between the entries present in both releases did not change: no close relation appeared or disappeared, and none changed type.

### 3.8 Sensitivity of the relation types to the thresholds

Changing one threshold at a time changed the classification of at most 15.5% of the 1,910,453 protein pairs examined (Table 10). Setting the identity threshold of the near-identical category to 95% or 99% moved 13.9% and 15.5% of the pairs, respectively, to a different type, and setting that of the same-family category to 85% or 95% moved 11.1% and 6.2%. Changing the coverage thresholds, by contrast, affected at most 0.7% of the pairs, so the classification is sensitive mainly to the sequence identity thresholds. In both cases the shift occurs between adjacent categories, near- identical and same family, or same family and homologue. Because the near-identical and length-variant categories share the same identity threshold, the size of the length-variant category also changes with the near-identical identity threshold (15,037 pairs at the default setting, 19,493 at 95% and 3,307 at 99%; Table 10).

**Table 10.** Sensitivity of the relation types to the thresholds. Classification of the pairs of distinct proteins above the search lower bound (1,910,453 pairs, for each pair the alignment direction with fewer differences) under the default setting (near-identical: *≥*98% identity and *≥*95% coverage; same family: *≥*90% and *≥*80%) and with one threshold changed at a time. The last column gives the share of pairs assigned to a different type than under the default setting.

| Setting | Near-identical | Length variant | Same family | Homologue | Changed |
|---|---|---|---|---|---|
| default | 358,789 | 15,037 | 384,586 | 1,152,041 | 0.0% |
| near-identical, identity ≥95% | 620,711 | 19,493 | 118,208 | 1,152,041 | 13.9% |
| near-identical, identity ≥99% | 74,600 | 3,307 | 680,505 | 1,152,041 | 15.5% |
| near-identical, coverage ≥90% | 372,913 | 922 | 384,577 | 1,152,041 | 0.7% |
| near-identical, coverage ≥99% | 353,611 | 19,515 | 385,286 | 1,152,041 | 0.3% |
| same family, identity ≥85% | 358,789 | 15,037 | 596,534 | 940,093 | 11.1% |
| same family, identity ≥95% | 358,789 | 15,037 | 266,388 | 1,270,239 | 6.2% |
| same family, coverage ≥70% | 358,789 | 15,086 | 384,633 | 1,151,945 | 0.0% |
| same family, coverage ≥90% | 358,789 | 14,120 | 383,955 | 1,153,589 | 0.1% |

Of the differences in the 373,826 close pairs, 75.2% are amino acid substitutions, 3.9% internal gaps and 20.9% unaligned ends. A total of 201 pairs (0.1%) differ only at their ends and are identical over the overlapping stretch, and of these 126 only at the N-terminus, which may indicate a different start codon annotation. As purely terminal differences are rare, no separate relation type was introduced for them.

### 3.9 Publication format

The complete mapping table, intended for checking, has 2,084,605 rows; in the normalised export, 427,581 protein pair relations remain. The reduction arises because the export stores every relation only once, per protein pair, and omits data that can be derived from the entries or other columns. The size thus decreases from 341.6 MB to 13.4 MB without loss of information, as the complete mapping table can be restored from the export.

### 3.10 Synthetic validation

For 99.6% of the 5,000 synthetic variants created from 500 curated proteins, the original protein was among the best hits, and for 99.0% it was the single best hit; the relation type agreed with the value expected from the modification for 99.5% of the variants and the difference count for 98.9% (Table 11). Most deviations from the expected value are not errors of the mapping. For CDSs without a start codon, if the first remaining codon is itself a possible start codon (e.g. GTG, TTG or ATT), translation converts it to methionine, so the N-terminus of the protein differs from the expected one. Whether this codon is a genuine start cannot be decided from the sequence, so these deviations arise from an inherent limitation of the method. Some C-terminally truncated variants were linked, as identical proteins, not to the original protein but to a shorter reference already present in the database, with which they are indeed identical. In a few cases, variants with amino acid substitutions became as close to another listed allele as to the original, and in one case the variant became identical to an already listed allele, which is to be expected in allele-rich families. In a few cases, the difference count was one higher than expected. In these, the substitution fell on the second position of the protein, and the local alignment also placed the initial methionine in the unaligned end. For one short protein (*nimC*), this also changed the classification: the variant became near-identical instead of same family, because identity is computed over the aligned stretch.

**Table 11.**
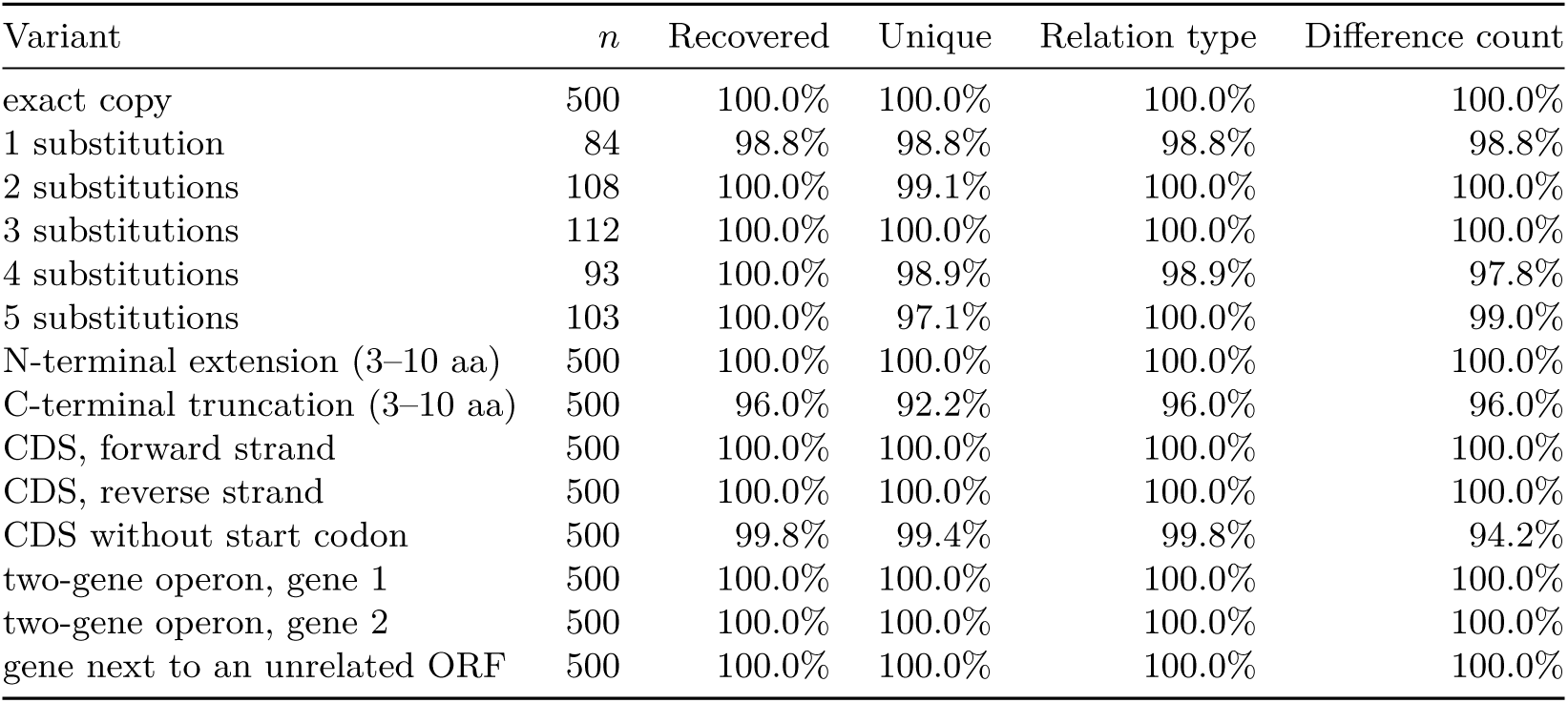
Synthetic validation. Recovery of variants created with known modifications from 500 randomly selected curated proteins of the release. Recovered: the original protein is among the best hits; unique: it is the single best hit; relation type and difference count: equal to the value expected from the modification.

### 3.11 Real-world use case

On the 100 genomes, AMRFinderPlus called resistance or biocide resistance genes at 1,110 loci, RGI at 3,609 and ResFinder at 793 loci. Locus-level agreement was 25.3% between AMRFinderPlus and RGI, 60.9% between AMRFinderPlus and ResFinder, and 19.0% between RGI and ResFinder; that is, the proportion of loci called by at least one of the two tools that were called by both. At the jointly called loci, the gene names agreed before mapping at 56.1%, 75.6% and 55.3% of the loci, respectively, whereas after mapping the references were identical or near-identical proteins at 93.3%, 94.2% and 89.6% (Fig. 9). At most of the shared loci, the tools thus reported the same protein under different names. Typical differences were the prefix (*blaTEM-1* and TEM-1), the species name in CARD names (*Staphylococcus aureus* LmrS and *lmrS*), spelling (*mph(A)* and *mphA*) and the variant letter of ResFinder (*blaTEM-1* and *blaTEM-1B*), but different allele numbers for the same protein also occurred (*aac(3)-IIe* and *aac(3)-IIa*). Every reference entry given by the tools was present in the release, so the relation of the two references could be determined at every jointly called locus.

**Figure 9.**
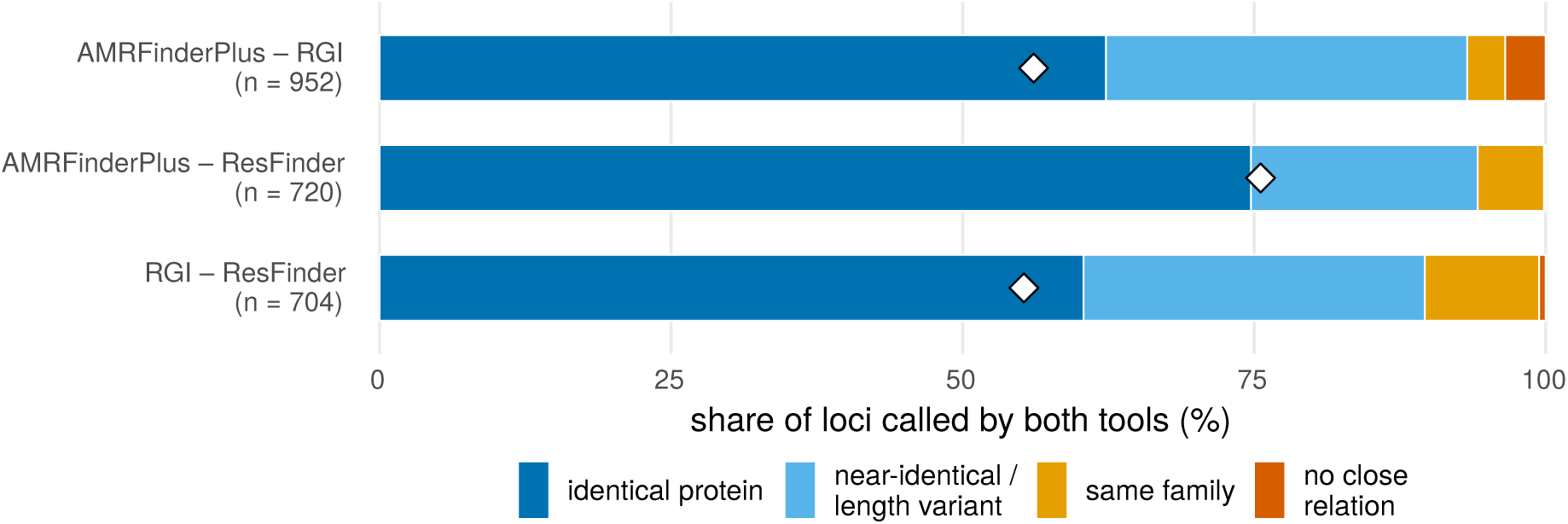
Agreement of the gene calls of the three annotation tools on 100 genomes, by tool pair, at the loci called by both tools. The bars show the relation of the references of the two tools in the release, and the diamond the name-based agreement before mapping. The number in parentheses after each tool pair is the number of loci called by both tools. No locus occurred at which a reference could not be linked to the release, so this category is not shown.

The differences remaining after mapping between AMRFinderPlus and ResFinder and between RGI and ResFinder were almost all different allele calls within the same family, accounting for 5.7% and 9.8% of the shared loci, respectively. Examples are *qacEdelta1* and *qacE*, *blaZ* and the PC1 *β*-lactamase of CARD, and *aac(6’)-Ib-cr* and AAC(6’)-Ib10. Between AMRFinderPlus and RGI, in addition, there was no close relation between the two references at 3.5% of the shared loci. At these, AMRFinderPlus reported the *Klebsiella kdeA* and RGI an *E. coli mdfA*-homologous efflux pump, or AMRFinderPlus the chromosomal *blaEC* and RGI the CMY-181 *β*-lactamase. At some of the shared loci (44 loci with AMRFinderPlus and 32 with RGI), ResFinder gave several equally good references, for example the *blaTEM-141*, *blaTEM-206* and *blaTEM-214* alleles for the same gene.

Locus-level agreement was limited mainly by the different content of the databases. For 69.6% (AMRFinderPlus and RGI), 38.6% (AMRFinderPlus and ResFinder) and 77.6% (RGI and ResFinder) of the loci called by at least one tool, the called reference has no close counterpart in the database of the other tool. RGI thus called chromosomal efflux pumps and regulators (e.g. *acrB*, *marA*, *emrR*) in every Gram-negative genome, AMRFinderPlus, compared with ResFinder, intrinsic resistance genes (e.g. *blaEC*, *emrD*, *mepA*), and ResFinder and DisinFinder genes not included in the other two databases, for example the *Salmonella aac(6’)-Iaa* gene, which is not classified in AMRFinderPlus, or the *sitABCD* operon. The share of loci whose reference would have had a close counterpart in the other database but which were called by only one tool was just 5.1%, 0.5% and 3.4%, respectively. Between AMRFinderPlus and RGI, about half of these were RGI hits to the *vanG* cluster.

## 4 Discussion

Our results show that name-based comparison of results obtained with different ARG databases is unreliable even for identical sequences. For 64.4% of the cross-database entry pairs encoding the same protein, the names differ even when letter case is ignored, and this share varies considerably between database pairs. Conversely, some identical names denote different proteins, and this also occurs for allele names. Although most differences arise from naming conventions and can be handled by rules, genuine conflicts can only be recognised by sequence. The comparison on real genomes showed the same. Where two annotation tools called the same locus, the gene names often differed, but after mapping the two references were mostly identical or near-identical proteins, and the remaining differences arose largely because the tools chose a different allele within the same family. Whether a locus is called by only one of the tools, however, depends mainly on the content of the databases. This cannot be resolved by name harmonisation, but the ARGeement release shows whether a missing call is due to the absence of a corresponding entry in the other database. We therefore see the standardisation proposed in our earlier review [9] as a sequence-based, versioned mapping rather than yet another nomenclature.

ARGeement differs from existing harmonisation solutions in that it links not the results of annotation but the entries of the databases themselves, with relation types expressing the closeness of the link (from identical protein to homologue) and with version tracking. hAMRonization [11] unifies the structure of the output of annotation tools, but does not map the gene names to each other, so the two tools complement each other. A gene list obtained with AMRFinderPlus in hAMRonization format can be translated with an ARGeement release into the nomenclature of CARD or ResFinder, together with the strength of the relation. Unlike argNorm [12], ARGeement does not normalise to an ontology but links the entries of the databases directly, with relation type and difference count, and follows the updates automatically. Unlike merged databases such as MEGARes [10], SARG [14], DeepARG-DB [15] and PanRes [16], it introduces no new nomenclature, so the names and curation of the source databases remain untouched.

The discrepancy between the uniform identity threshold and the curated family boundaries is not a methodological error, but reflects the difference between the family concepts. Representative-based clustering excludes chaining, so the remaining discrepancies arise because the curated family boundaries do not follow a single identity threshold. The *Klebsiella β*-lactamases illustrate this well. The boundaries of the SHV, LEN and OKP families are tied largely to the host species (*K. pneumoniae*, *K. variicola*, *K. quasipneumoniae*), which is not reflected in protein sequence identity alone, so the clusters obtained with a single identity threshold do not coincide with these families: some families merge into a common cluster, others are split across several clusters. For family-level comparisons, the curated NCBI family label is therefore more reliable, whereas the sequence-based cluster reflects only the proximity of the sequences.

The family concepts of CARD and NCBI can mostly be nested, with the two systems describing the same gene group at different levels. Where, however, the NCBI hierarchy does not group the members of a CARD family under a common node, the two systems are organised by different criteria. The vancomycin resistance genes illustrate this well. CARD places the Van ligases in one family by function and forms a separate family from the complete glycopeptide resistance gene cluster, whereas the NCBI hierarchy divides the ligases into two branches by their product (D-Ala-D-Lac ligases, e.g. VanA and VanB, and D-Ala-D-Ser ligases, e.g. VanC and VanE), without a common node. One system regards the enzyme function as primary, the other the type of resistance, and neither is more correct than the other. For family-level comparisons between databases, it must therefore always be stated which family concept is used.

The harmonisation of drug classes leads to a similar conclusion. Assignment to common classes necessarily loses the finer subdivisions, and some native labels do not denote a separate resistance class at all. The CARD labels azole antibiotic and imidazole antibiotic, for example, belong exclusively to CTX-M *β*-lactamases, and their mechanical assignment would have overestimated the frequency of nitroimidazole resistance many times over. Comparing the classifications of the databases therefore requires expert review, as we also emphasised in our earlier review [9].

Database updates also affect the interpretation of earlier analyses. With an earlier database version, a gene whose allele was added only later can be identified at best as a close allele or at the family level, if at all. Pinned profiles and the comparison of releases show which entries are affected, so the affected genes can be checked specifically in a previously analysed sample set, without repeating the whole analysis. Sequence-based comparison can also point to errors in the reference sequences. In two ResFinder entries (*mph(D)*, *blaFLC-1*), the gene cannot be read in full length because of an internal stop codon or a length not divisible by three, and none of their ORFs is linked to a curated protein, which indicates a frameshift in the reference sequence.

The tool has several limitations. Mutation-based determinants are left out of the mapping. Sequence identity does not guarantee functional identity, and in allele-rich families ‘near-identical’ denotes the closest, but a different, allele. The relation types are based on fixed, relative thresholds, which do not fit the curated boundaries in every family. As the thresholds are given as proportions, the same number of differences may fall into a different type for a short protein than for a long one. Five amino acid substitutions, for example, remain near-identical at an identity threshold of 98% only for proteins of at least 250 amino acids. In ORF splitting, the choice of start codon carries an uncertainty of a few amino acids, and genes shorter than 60 amino acids or truncated at the boundary of the entry may be missed. For ORFs of genes absent from the curated databases, the selection is based only on the covered length. The family clusters also depend on the choice of representatives. Choosing the longest protein as representative is deterministic, but the appearance of a new, longer entry after an update may rearrange the clusters. The derived family labels rely on the curation of AMRFinderPlus, so genes present only in other databases have no family label. Drug class harmonisation is based on a manually maintained table, whose gaps are reported for every release.

Possible directions for further development are the position-based comparison of mutations using the reference sequences and the inclusion of results annotated with aggregating databases (MEGARes, SARG, PanRes). The entries of the aggregating databases derive largely from the same primary databases, so they can be assigned to the entries of a release by their sequence, and the results obtained with them can thus also be translated to the names of AMRFinderPlus, CARD or ResFinder.

## 5 Conclusions

ARGeement maps the entries of three major ARG databases to each other on a sequence basis, with version tracking, and produces a documented, publishable result for every database release. On 100 real genomes, mapping raised the agreement of three annotation tools at shared loci from 55.3%–75.6% (by name) to 89.6%–94.2% (by protein), whereas whether a gene was called at all depended mainly on database content. The results show that name-based comparison between databases can lead to erroneous conclusions even for identical sequences, and that sequence-based mapping and curated family definitions complement each other. When reporting the results of ARG analyses, it is therefore advisable to report the exact database version alongside the gene names, and to use a sequence-based mapping for comparing results obtained with different databases.

## Acknowledgements

In memory of Péter Gálfid, to whom we owe the establishment of the Centre for Bioinformatics.

## Author contributions (CRediT)

*AGT* : Review & Editing; *IC* : Funding acquisition, Review & Editing; *GH* : Formal analysis, Review & Editing; *NS* : Conceptualization, Formal analysis, Methodology, Review & Editing, Software, Supervision, Validation, Visualization, Writing – original draft; *VV* : Formal analysis, Review & Editing, Writing – original draft. All authors read and approved the final manuscript.

## Code availability

The source code of ARGeement is available under the MIT licence: https://github.com/solymosin/ARGeement. The database releases (the mapping tables, the schema description and the content report) are deposited in a versioned Zenodo record under the CC BY 4.0 licence; the release described in this article (2026-11-16_184628) has the DOI 10.5281/zenodo.24300987, and the concept DOI always pointing to the latest release is 10.5281/ zenodo.24278768. The release of the pinned profile amrfinder-2026-08-07 used in the comparison of database versions (2026-11-16_203603) is deposited in a separate Zenodo record (DOI 10.5281/zenodo.24301243; concept DOI 10.5281/zenodo.24301242). A summary of the latest release is available at https://solymosin.github.io/ARGeement/. From CARD, the package contains only the ARO accessions and terms (CC BY 4.0), and no sequence from any source database; the use of the source databases is subject to their own terms. The accession numbers of the genomes used in the real-world use case and the locus-level results are in the concordance/ directory of the GitHub repository.

## Ethics statement

Not applicable. The study used only publicly available database records and genome sequences; it involved no human participants or animals.

